# Dynamic characteristics rather than static hubs are important in biological networks

**DOI:** 10.1101/2020.09.30.320259

**Authors:** Silke D. Kühlwein, Nensi Ikonomi, Julian D. Schwab, Johann M. Kraus, K. Lenhard Rudolph, Astrid S. Pfister, Rainer Schuler, Michael Kühl, Hans A. Kestler

## Abstract

Biological processes are rarely a consequence of single protein interactions but rather of complex regulatory networks. However, interaction graphs cannot adequately capture temporal changes. Among models that investigate dynamics, Boolean network models can approximate simple features of interaction graphs integrating also dynamics. Nevertheless, dynamic analyses are time-consuming and with growing number of nodes may become infeasible. Therefore, we set up a method to identify minimal sets of nodes able to determine network dynamics. This approach is able to depict dynamics without calculating exhaustively the complete network dynamics. Applying it to a variety of biological networks, we identified small sets of nodes sufficient to determine the dynamic behavior of the whole system. Further characterization of these sets showed that the majority of dynamic decision-makers were not static hubs. Our work suggests a paradigm shift unraveling a new class of nodes different from static hubs and able to determine network dynamics.

## Introduction

Recently, there has been a shift in biological research from studying single compounds towards complex regulatory networks and their dynamic behavior (Kitano, 2002). Hence, network analysis has emerged as a powerful tool to understand complex biological processes (Barabási et al., 2011). In accordance, diseases or the development of cancers are rarely a consequence of a mutation of a single component within a network, but rather its perturbation (Barabási et al., 2011; Barabási and Oltvai, 2004). Thus, to understand the development of diseases, we have to follow the network dynamics (Sverchkov and Craven, 2017). However, a significant limitation is that interaction graphs cannot adequately represent temporal changes of a pathway or network. Among models that investigate dynamics, Boolean network models (Kauffman, 1969) can approximate simple features of interaction graphs integrating also dynamics. Here, known regulatory interactions can be directly formulated as Boolean functions (*4*)(Naldi et al., 2015) which can then be used to obtain the dynamics of the network. The dynamic simulation of a Boolean network leads to periodic sequences of states, called attractors, describing the long-term behavior of the model (Xiao, 2009). Attractors depict the activity of each component within the system at a specific point in time and can be associated with phenotypes (Kauffman, 1993). In this sense, the sequence of attractor states and the compounds involved can be seen as a model of the activities specific for a biological state.

Biological networks exhibit a scale-free topology (Barabási and Oltvai, 2004). That is, a small number of components called hubs are highly connected, whereas the majority of nodes have few connections (Guimerà and Nunes Amaral, 2005). Often hubs are seen as the master regulators of biological processes (Borneman et al., 2006; He and Zhang, 2006) whose removal correlates with lethal phenotypes (Jeong et al., 2001). However, studies state that it is almost impossible to eliminate the activity of hub nodes by targeted inhibition with small molecules (Lu et al., 2007; Song et al., 2017). The most striking difference between both observations is the fact that one side considers the complete loss of proteins while the other side thinks about inhibition. Again, these studies are mainly based on static interaction graphs. Since master regulators are supposed to affect the behavior in time of biological processes, it is appealing to investigate them from a perspective involving network dynamics. Nevertheless, investigating and detecting master regulators by their impact on network dynamics comes with the limit given by the complexity of the system. This might lead to unfeasibility of dynamic investigation or to reduction procedures, that might in turn alter the dynamic behavior. Hence, strategies to bridge the gap needed to unravel the dynamics of biological networks are of crucial interest. Which nodes are then crucial for the dynamics of a biological network? How can they be detected? Are these dynamic regulators actually the already widely studied static hubs? Can these nodes be relevant as intervention targets? To uncover these questions, we studied the dynamics in biological systems (Figure 1).

**Figure 1.**
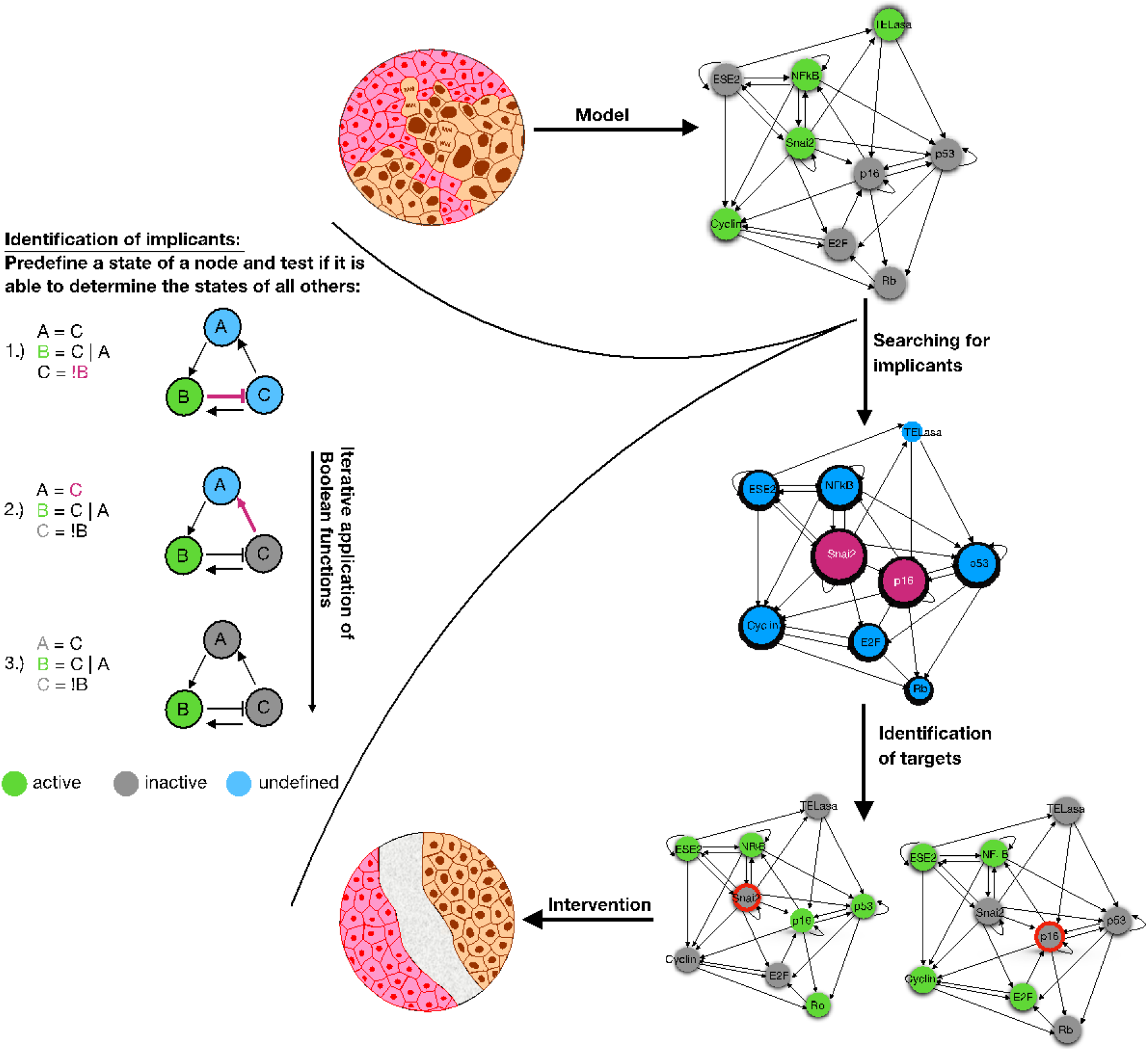
Scheme of the implicant-analysis. A biological system is transferred to a Boolean network model (epithelial to mesenchymal transition model (Méndez-López et al., 2017), upper interaction graph on the right). Next, this model is used to identify dynamic determining nodes (here called implicants, purple nodes in the right graph, middle). Perturbations of these potential targets are tested in silico (lower-right graphs) and promising targets are transferred to intervention screenings. The left part of the figure demonstrates the effect of an implicant in a small illustrative example. In step 1, implicant B is set to 1 (green node). Subsequent application of the Boolean functions (purple arrows) determines the value of C (0, gray node, step 2) and A (0, gray node, step 3). Consequently, the assignments of all nodes are determined by implicant B.

Here, we present techniques on how to identify nodes sufficient to determine the dynamic behavior of the network and discuss the implications of targeting these nodes for intervention experiments

## Results

### Small sets of compounds determine the dynamics

The state of a Boolean network is given by a vector of values (0/1) assigned to their nodes (n). A state change is a result of the application of a Boolean function for each node (Kauffman, 1969). The state change of a node thus depends on the previous state of other nodes. We assume that not all nodes are necessarily involved in the dynamic behavior of the network. Therefore, we introduce a group of nodes, called implicant sets, from which it is possible to determine a state of the system and thus its dynamics for every assignment of values by iteratively applying Boolean functions.

To confirm our hypothesis, we implemented two strategies to search for a minimal set of implicants (k) in selected biological representative and scale-free Boolean network models (Appendix 1–Algorithm 1 and Appendix 1—Algorithm 2): an exhaustive search and a heuristic search. Since the search space is 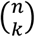 and as a consequence the computation time grows exponentially with each additional component of the implicant set, exhaustive search is feasible only for small networks (Hopfensitz et al., 2013). Thus, we applied the full search only as a reference benchmark to evaluate our newly established heuristic search algorithm. Additionally, we removed redundant nodes from the network to improve the running time of both algorithms (Appendix 1—Algorithm 3).

Finally, we compared the performance of both algorithms. The heuristic defined a minimal set of implicants in 54.3% of the networks. Furthermore, in 32 out of the 35 networks considered, the implicant set defined by the heuristic was a superset of a minimal implicant set found by exhaustive search. The performance of the heuristic compares quite favourably considering the required running times of 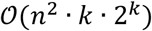 for the heuristic search and 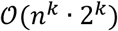 for the exhaustive search to calculate an implicant set of size *k* of a network with *n* nodes. Furthermore, our analyses revealed that the typical size of minimal implicant sets is small (Figure 2A). The cardinality of implicants ranges from 1 to 9 for an average size of 19 nodes per network. Here, the exhaustive approach identified a mean of 4.4 implicant nodes and the heuristic a mean number of 5.1 implicant nodes. Only a low correlation between the network size and the size of the implicant set could be found (Pearson correlation at 0.32 using the exhaustive strategy and 0.39 using the heuristic strategy). Furthermore, the relation between the set size of the implicant set and the network size is not linear (*R^2^*= 0.104 using the exhaustive strategy and *R*^2^ =0.153 using heuristic strategy).

**Figure 2.**
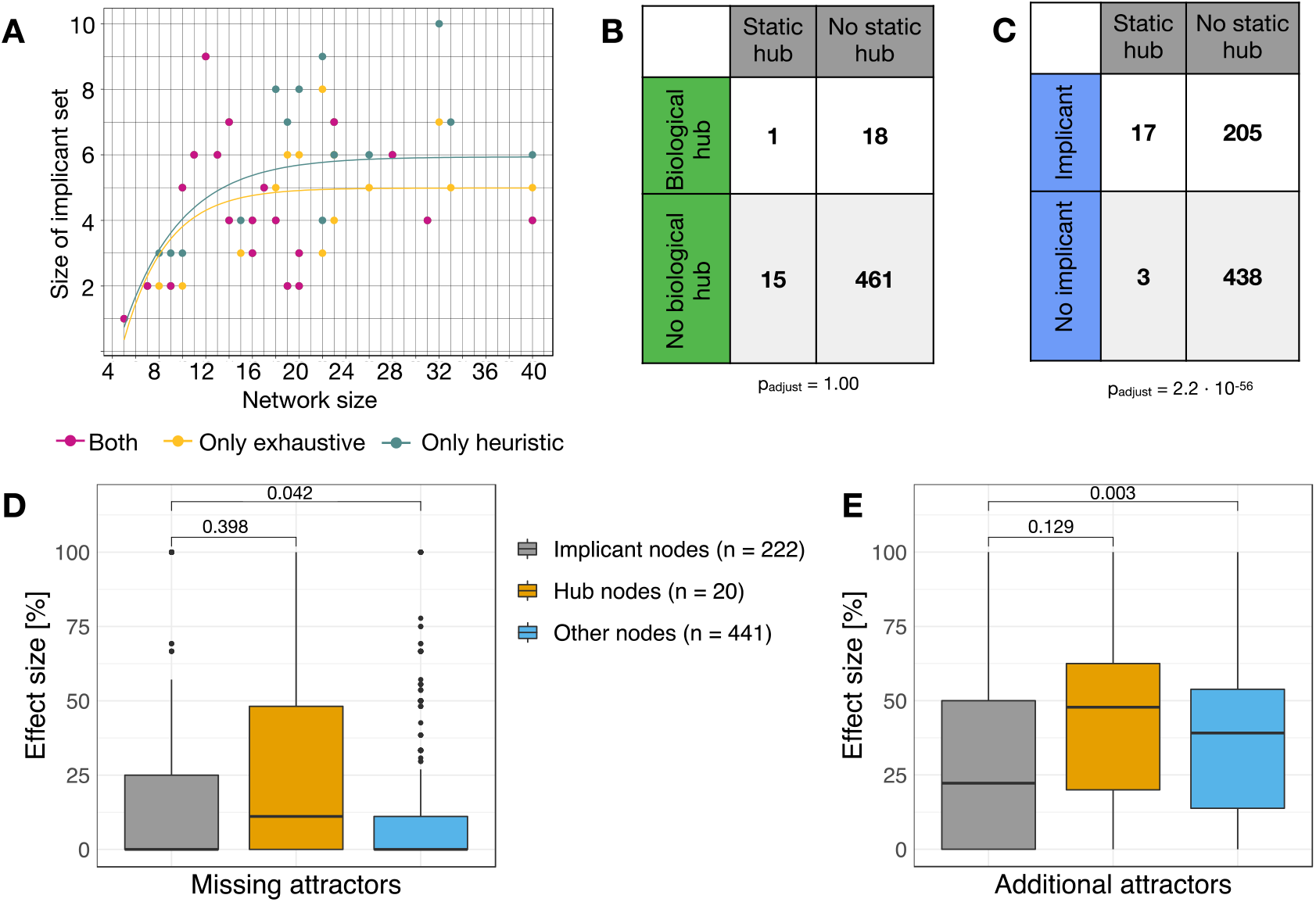
Dynamic in silico analyses. (A) The implicant set is a set of nodes whose determined states are enough to determine the entire dynamics of a network by applying iteratively Boolean functions. The overall size of the implicant set is small. Two strategies were employed to search for minimal implicant sets. The exhaustive search (yellow) identifies a minimal set by checking all possible combinations. In contrast, the heuristic (grey) assesses the importance of components. The size of implicant sets that were identified by both search approaches is given in pink. The relation between the size of the implicant set to the network size is non-linear (logarithmic fit). (B) Regulatory interactions of nodes in Boolean network models are comparable to their biological representatives. (C) Majority of implicants are no hub nodes. Nodes are defined as hubs if their z-score was >2.5 (Guimerà and Nunes Amaral, 2005). Statistics were performed with Cochran’s Q test with post-hoc pairwise sign test with Bonferroni correction. (D) Percentage of missing attractors or (E) additional attractors (side effects) after interventions (knockouts/overexpression; Wilcoxon test). We assume significant results if p < 0.05.

### Majority of implicants are lowly connected but behave like hubs

Next, we studied the connectedness of the dynamic influencing nodes of the implicant sets to uncover if they are hub nodes. To ensure that the links of nodes within the considered Boolean network models are comparable to their biological representatives, we initially compared them to the interactions of their representatives found on BioGRID (Stark et al., 2006). Here, we did not find any differences (p = 1.0), concluding that the static interactions in the Boolean network models are comparable to biological interactions (Figure 2B).

Across all considered Boolean network models, only 3.2% of compounds could be identified as static hubs (z-score >2.5 (Guimerà and Nunes Amaral, 2005)). Comparing these nodes with our implicant sets showed that they differ significantly (p = 2.2·10^−56^) (Figure 2C).

Based on these results, we analyzed the effect of targeted interventions, i.e. permanent overexpression or knockout on implicants, static network hubs or other nodes (Figures 2D and 2E). Here, we focused on the case if interventions have an impact on the number of missing attractors in order to estimate the effect size or whether the interventions lead to additional attractors and thus side effects. Interestingly, implicant nodes perform comparable to hub nodes in both analyses but significantly better than other nodes. Also, interventions with implicant nodes evoke fewer side effects than interventions with hub nodes.

Hence, targeted interventions of dynamic influencing nodes appear to be a new possibility to achieve a sufficient effect size under a reduced risk of side effects.

### Interventions of lowly connected implicants can change phenotypical behavior

Until now, we analyzed the general dynamics within our considered Boolean network models. However, a model is only adequate if it can reflect reality. To estimate if our implicant nodes also play a role in the dynamics of real biological processes and are not artefacts of the modeling approach, we compared the simulation outcomes of interventions with lowly connected implicants with the findings from biological experiments.

Cohen et al. (Cohen et al., 2015) describe molecular pathways of tumor development to invasion and metastases. In their network are two statically and biologically lowly connected implicants, AKT2 (z-scorestatic= 0.98; z-scoreBioGRID= −0.33) and TWIST1 (z-scorestatic= 0.04; z-scoreBioGRID= −0.36). While the simulation of AKT2 overexpression reached attractors supporting tumor development by inhibiting apoptosis and activation of epithelial to mesenchymal transition (EMT), in silico knockout of TWIST1 prevents tumor associated characteristics. Laboratory experiments support both simulations. For AKT2, they describe that it mediates EMT by inhibiting GSK3*β*/Snail signaling (Lan et al., 2014, p. 2) and that its overexpression in combination with PTEN loss promote metastases (Rychahou et al., 2008). Both events thus support tumor formation. In contrast to the negative effect of AKT2, the favourable effect of TWIST1 could also be confirmed. Here, a knockout of TWIST1 in breast cancer cells inhibited the expression of EMT markers and prevented metastases in immune-deficient mice (Xu et al., 2017).

Another example of the biological impact of lowly connected implicants can be found in the network of Méndez-López et al. (Méndez-López et al., 2017) also dealing with EMT. All implicants (Snai2, ESE2 and p16) within this network are statically (z-score_static_: 1.46; 0; 0.97) and biologically (z-score_BioGRID_: −0.5; −0.59; 0.66) lowly connected. The strongest intervention effect can be observed by targeting the implicant node Snai2. While the unperturbed network simulation ends in three single state attractors representing epithelial, senescent and mesenchymal characteristics (Méndez-López et al., 2017), the simulation of Snai2 overexpression only yields one attractor with mesenchymal characteristics (Méndez-López et al., 2017). The attractor with mesenchymal characteristics disappeared by simulating Snai2 knockout (Méndez-López et al., 2017). Laboratory experiments also support these effects of Snai2 in the Boolean network model. Here, in vitro overexpression of Snai2 resulted in a mesenchymal appearance of cells within 72 hours (Chakrabarti et al., 2012) while depletion of Snai2 supports premature differentiation (Mistry et al., 2014).

Lowly connected implicant nodes influencing dynamic behavior cannot only be found in Boolean network models associated with cancer. The Boolean network of Krumsiek et al. (Krumsiek et al., 2011) describes haematopoiesis. Based on our analysis, we identified six implicant nodes which are statically and biologically lowly connected. A knockout of each of these proteins leads to the loss of a blood cell lineage in the simulation, while abnormal states are absent (Krumsiek et al., 2011). This is in line with results from in vitro experiments (Karsunky et al., 2002; Kawada et al., 2001; Laslo et al., 2006).

To sum up, literature comparison of our simulations of interventions with implicants could confirm our results independent of the cell context. Based on these results, it can be reasoned that even nodes with only a few connections in a network structure can change the phenotype of biological processes. Our new method helps to identify exactly these dynamically important nodes.

### Implicant screening identifies promising candidates for treating colorectal cancer patients

Based on an extensive literature search, we could already show that our method is capable of identifying promising intervention targets. In the next step, we now guide laboratory experiments based on a modelling approach and the analysis of its dynamics. For this purpose, we developed a new Boolean network model (Appendix 2—Table 1) dealing with the crosstalk of two frequently mutated pathways in colorectal cancer – Wnt and MAPK (Jeong et al., 2018)- with the focus on KRAS driven tumors.

Our method identified seven implicants in this newly established Boolean network model. Two of them, ERK and TCF/LEF are static network hubs, while the other five (APC, CIP2A, AKT1, RAC1 and GSK3*β*_deg) are lowly connected components (Figure 3A).

**Figure 3.**
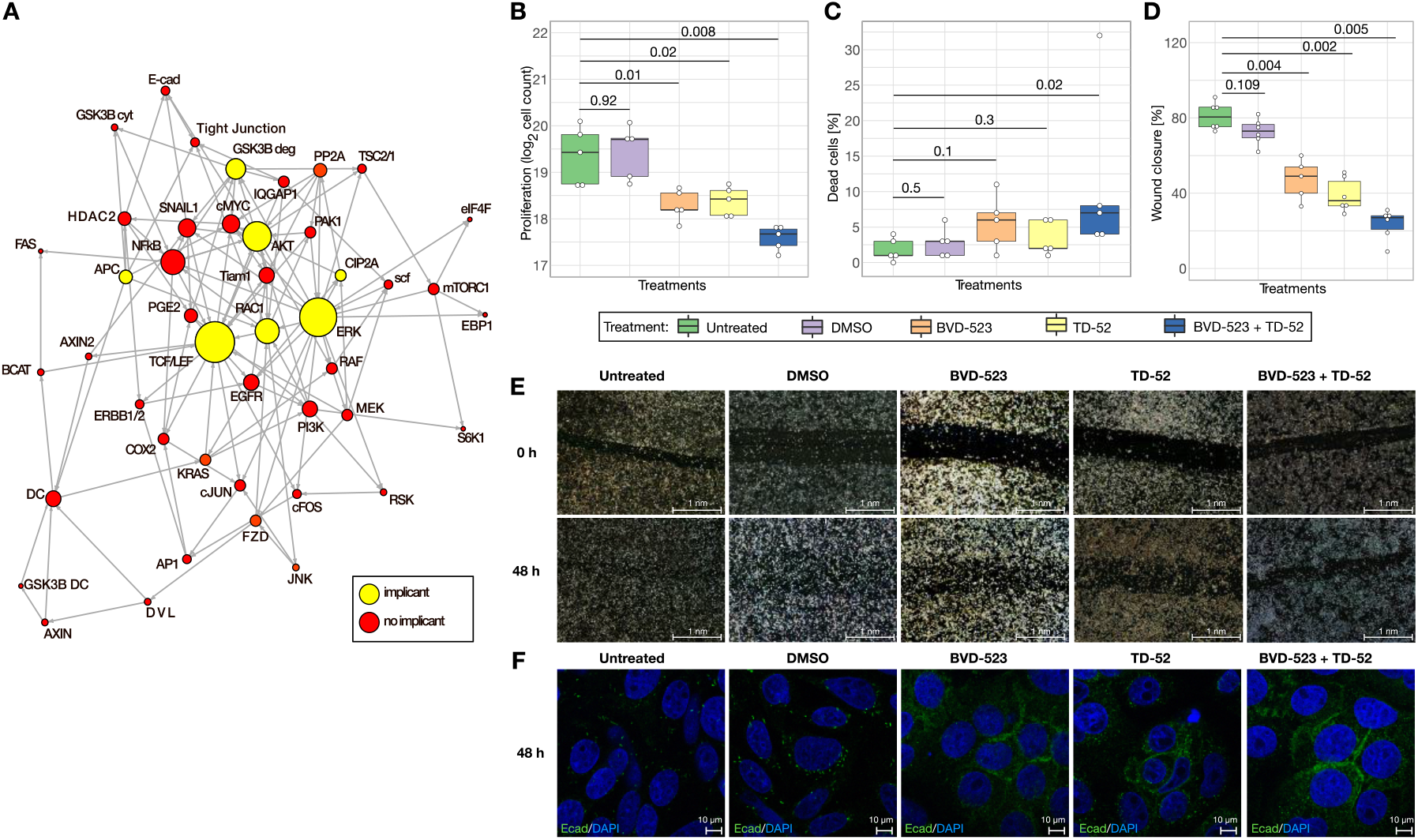
Targeting implicants in vitro. ERK and CIP2A identified as implicants were targeted individually and in combination. (A) Interaction graph of the colorectal cancer model. The circle size gives the number of regulatory interactions; implicant nodes were identified by the heuristic approach (yellow). (B) Cell counts from proliferation assay after 24 hours post-treatment (n=5, Wilcoxon test). For both of the single drug treatments a 2-fold reduction was detected. The combined treatment led to a 3.4-fold decrease. (C) Percentage of dead cells from proliferation assay shows no significant differences in apoptosis (n=5, Wilcoxon test). (D, E) Percentage of wound closure after 48 hours post-treatment indicates a reduced migratory potential (BVD-523: n=5, otherwise: n=6, Wilcoxon test). (F) Merged confocal microscope pictures of E-cadherin staining (green) and colored nuclei (blue) 48 hours post-treatment. Treatment of implicants restored E-cadherin at the cell membrane.

Based on the availability of small molecules for human treatment, the requirement that the target has not been tested in clinical trials for colorectal cancer patients and that no resistances are known for the inhibitor, we selected the most promising intervention candidates out of the list of implicants and studied their effect. Loss of functions of the implicant APC is one of the most frequently occurring alterations in colorectal cancers (Kinzler and Vogelstein, 1996). However, there are still no treatment options for APC loss in humans. Taking this into account, we have chosen the colorectal cancer cell line SW480 for our experiments, which expresses a truncated form of APC (Bienz and Hamada, 2004). After intensive literature search and intervention simulation (Appendix 3), we decided, as a first step, to investigate the impact of the less connected component CIP2A. This target alone already shows promising intervention effects and has not been evaluated in the context of colorectal cancer.

Nevertheless, we also investigated the possibility to suggest a coupled intervention by further considering the hub ERK. For this purpose, we selected the specific ERK inhibitor BVD-523 (Germann et al., 2017) and the specific CIP2A inhibitor TD-52 (Yu et al., 2014). APC loss is associated with loss of cell adhesion (Bienz and Hamada, 2004), as well as increased proliferation (Heinen et al., 2002) and migration (Kawasaki et al., 2003). Therefore, we studied the influence of our interventions on these effects.

Treatment with either BVD-523 or with TD-52 reduced the proliferative potential of SW480 by 2-fold (mean values, Figure 3B) within 24 hours in comparison to untreated or DMSO treated controls without increasing apoptosis (Figure 3C). By combining both approaches of inhibition, an even stronger mean inhibitory effect of 3.4-fold was achieved (Figure 3B). In addition, the migratory potential of ERK or CIP2A treated SW480 cells was significantly reduced (Figures 3D and 3E) and E-cadherin was restored at the cell membrane (Figure 3F and Appendix 4—Figure 1) thereby supporting that cell adhesion further reduces the migratory potential. Moreover, these effects are further enhanced when the treatment is combined.

These initial in vitro experiments, which were performed based on implicants identified in the network structure of a Boolean network model, already show significant results in the inhibition of proliferation and migration of colorectal cancer cells — leading to the conclusion that the consideration of dynamic characteristics has an added value for the identification of promising therapeutic targets. Thus, we presume that dynamic characteristics should be given more significant consideration and should be taken into account when planning interventions.

## Discussion

Network approaches are already being used to identify disease modules. For example, Goh et al. (Goh et al., 2007) studied the connection of diseases and involved genes, while Wang et al. (Wang et al., 2013) studied the chance of a drug to pass clinical trials based on the connection of its targets.

In this study we used Boolean network models. These simple models capture the complex biological behavior from homeostatic processes (Fauré et al., 2006; Ikonomi et al., 2020) to cancer signaling (Cohen et al., 2015; Dahlhaus et al., 2016; Méndez-López et al., 2017) or developmental processes (Krumsiek et al., 2011; Siegle et al., 2018). Moreover, they can be generated from qualitative knowledge about interactions (Naldi et al., 2015). Additionally, it is possible to create Boolean network models from time-series data (Schwab et al., 2017). Here, we recommend identifying promising candidates for therapeutic approaches employing dynamic characteristics of the biological processes involved. For this purpose, we present a method which makes it possible to identify dynamic influencing nodes in the wiring of biological networks in a reasonable time.

Exhaustively searching for implicants can be time-consuming if not even impossible for larger Boolean network models. To ease this limitation, we established a heuristic search procedure. In order to validate our new approach, we analyzed our chosen Boolean networks also by exhaustively searching for implicants. A comparison of the required time already shows that the heuristic approach was much faster. Altogether the heuristic found an implicant set up to 1 million times faster than the exhaustive search. Nevertheless, it should be kept in mind that it gives not always the minimal implicant set. 91.4% of the implicant sets determined by the heuristic were equal to or a superset of the minimal implicant set determined by the exhaustive search, indicating an excellent performance of the heuristic. Furthermore, redundant nodes in the supersets of implicants can be identified by additional simulations.

Many previous approaches working with networks perform static analyses and developed methods to identify important nodes (Zhu et al., 2007). In contrast to them, we chose a dynamic approach that incorporates temporal changes of a compound together with its long-term behavior within the complete network. In fact, it is shown that only studying interaction mechanisms from a static point of view can result in insufficient characterization of the final behavior. Exemplarily, inhibitory studies with small molecules showed reactivation of silenced kinases due to reactivation mechanisms (Song et al., 2017). Hence, studying overlap of static interactions essentially is not enough to understand complex dynamics leading to reactivation and resistance mechanisms. Altogether supporting the idea of looking for suggestions emerging from the dynamic analysis of these networks.

Thus, we can assess the impact of a compound based on phenotypical changes. With our approach, we differentiate ourselves from other dynamic studies. General studies by others suggests that hub compounds are master regulators of biological processes (Jeong et al., 2001). In contrast, it could be shown that only 20% of hubs are essential genes (Goh et al., 2007). Being aware of the fact that the definition of essentiality is context dependent, we quantified the proportion of essential genes being implicant or hubs in our set of networks. By screening the HEGIAP database (Chen et al., 2019), we found 12% of our compounds classified as essential in humans. Of these 27.5% are implicants and only 4% are hubs. Moreover, all essential hubs identified were also implicants. Hence, supporting the finding that implicants are essential gatekeepers of dynamics. Again, other studies postulated different subgroups of hubs by estimating their interactions over time from compilations of yeast mRNA expression profiles and static protein-protein interaction graphs. Based on this data, there is indication of the impact of time and space and relevance of two different classes of hubs. One group of compounds which interact with their partners simultaneously and another group which interacts at different times and/or locations (Han et al., 2004). Complimentary to that, our approach – based on dynamic models – not only shows the different impact of static hubs but also reveals another set of compounds which determines the dynamics of an interaction network. We could show that a small set of components is sufficient to determine the dynamics of a complete network. We called these nodes implicant set. This finding is consistent with the observation that alterations of individual proteins can disrupt complex signaling cascades, ultimately supporting the development of diseases and cancer (Barabási et al., 2011; Barabási and Oltvai, 2004).

One such protein of the implicant set is APC. APC is seen as an initiator of colorectal cancer, which is mutated in 80-85% of cases (Kinzler and Vogelstein, 1996). Note that, besides being one of the first inducers of colorectal cancer, APC seems to be an attractive therapeutic target. Zhang et al. (Zhang et al., 2016) presented in first preclinical studies, a small molecule called TASIN-1 (truncated APC selective inhibitor) that selectively eliminates cells with truncated APC by a mechanism that bases on synthetic lethality (Zhang et al., 2016). The fact that APC was identified as an implicant in the Boolean network model of Wnt and MAPK signaling in colorectal cancer supports its validity. Also, further implicants could be identified in this model whose in vitro inhibitions resulted in a reduction of tumor characteristics and thus phenotypical changes. Comparable to APC, there are only preclinical studies with TCF/LEF inhibitors as well as RAC1 inhibitors which are all not yet tested for humans. Contrary, AKT inhibitors have already shown insurgence of resistance in colorectal cancer patients (Song et al., 2019) while a controversial role of GSK3*β* is still under discussion (Shakoori et al., 2005). These results lead to the reasoning that the analysis of network dynamics can identify disease nodes as well as promising intervention targets.

Besides studying the impact of interventions on single implicant nodes, it is also possible to study their combinations. Having a small set of nodes to combine makes the search for promising combinations much easier and faster.

To propose a drug intervention based on our in silico results, we selected the two inhibitors BVD-523 (Germann et al., 2017) and TD-52 (Yu et al., 2014) to test the impact on inhibiting our two implicants ERK and CIP2A. Besides phenotypical attractor evaluation, the choice was made because both inhibitors are already being tested for other cancers than KRAS driven colorectal cancer in human clinical trials (https://clinicaltrials.gov, 9 studies for BVD-523 (NCT03417739, NCT02994732, NCT02296242, NCT01781429, NCT03454035, NCT03698994, NCT02608229, NCT02465060, NCT03155620) and TD-52 a derivative of erlotinib, a drug already approved for treatment of lung tumors (830 studies on http://clinicaltrials.gov)). Also, it is known that both inhibitors are specific, and their mechanisms of action are known (Germann et al., 2017; Yu et al., 2014). Initial laboratory experiments showed good results for both implicants in reducing tumor characteristics. Nevertheless, a combination of the hub ERK with the less connected CIP2A yields a cumulative effect.

Coupling kinase and phosphatase inhibitors have already been suggested in literature to prevent known insurgence of resistance of MEK and RAF inhibitors (Westermarck, 2018). Furthermore, this idea is supported by findings showing that PP2A status impacts the response to MAPK inhibitors (Kauko et al., 2018). The inhibition of our two chosen implicants has the ultimate goal of reducing the activity of ERK. While BVD-523 blocks ERK activity by competing for ATP binding(Germann et al., 2017), TD-52 acts by blocking CIP2A promoter binding. Thereby CIP2A inactivation enhances the activity of PP2A, which can inhibit ERK phosphorylation (Yu et al., 2014). Based on this regulation, we perhaps circumvent a reactivation mechanism by treating different activation mechanisms of ERK as described by Song et al. (Song et al., 2017).

In this paper, we introduce the notion of implicants, defining a set of components that can determine the dynamics of a whole network. Intervention studies with these components support the idea that they are promising candidates for therapeutic approaches. Thus, we believe that dynamic characteristics rather than static hubs are essential factors to be considered when developing drugs or planning laboratory intervention studies. With our approach, we approximate personalized medicine by creating models that are individually adapted to the patient and identify the best possible intervention candidates through simulation.

## Materials and Methods

### Boolean networks

For the definition of Boolean networks (Kauffman, 1993, 1969) and established attractor search algorithms see e.g. the BoolNet (Hopfensitz et al., 2013; Müssel et al., 2010, p.) package. We briefly recall the most important notations.

Boolean variables 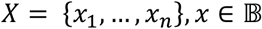 represent compounds with a state of 1 (expressed/active) or 0 (not expressed/inactive). Biochemical reactions are represented by Boolean functions 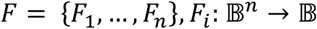 (Kauffman, 1993, 1969). If the value of Fi depends on *x*_*i*1_, *x*_*i*2_, …., *x*_*ik*_ let *f_i_* denote the function defined on these inputs, i.e. *F_i_*(*x*) = *f_i_*(*x*_*i*1_, *x*_*i*2_, …, *x_ik_*). *f_i_* is also called the Boolean function for the i-th position (e.g. gene) of F. To analyze the dynamics of Boolean networks over time, the state of the networks *x*(*t*) = (*x*_1_(*t*), …, *x_n_*(*t*)) is defined by the states of all variables *x_i_* at point in time *t*. In synchronous Boolean network models, all Boolean functions are updated at the same time to proceed from state at time t to its successor state at time t + 1 which is defined by *x*(*t* +1) = (*F*_1_(*x*(*t*)), …, *F_n_*(*x*(*t*))) of x(t). The dynamics of Boolean network models can be viewed in a state transition graph linking each state of the state space to its successor state. The state-space of Boolean networks with *n* nodes is finite with 2*^n^* possible states. Thus, the model will eventually enter a recurrent sequence of states called attractors (cycles) depicting the long-term behavior of the network. In a biological context, attractors are associated with phenotypes.

All Boolean network simulations were performed with R v3.4.4 (R Core Team, n.d.) and the R-package BoolNet (Müssel et al., 2010) v2.1.5.

### Boolean network model selection

For our analysis we extracted Boolean functions of Boolean network models from pubMed with the search item “Boolean network model” as well as Boolean functions from the Interactive modelling of Biological Networks database (https://cellcollective.org). Networks were selected until May 24^th^ 2017. We excluded networks whose dynamics could not be investigated in feasible time, networks with too many attractors and networks in which nearly all nodes are input nodes. Additionally, we excluded non-scale-free networks and those which could be reduced up to their input nodes. In total, we analyzed 35 networks with a size ranging from 5 to 40 nodes (Table 1).

**Table 1.** Boolean network models investigated. Depicted are the described process as well as the organism for which the model was setup.

| Network | Process | Organism |
| --- | --- | --- |
| (Azpeitia et al., 2013) | Root stem cell niche | *Arabidopsis thaliana* |
| (Brandon et al., 2015) | Oxidative stress response | *Aspergillus fumigatus* |
| (Calzone et al., 2010) | Cell-fate decision | *Homo sapiens* |
| (Cohen et al., 2015) | EMT | *Homo sapiens* |
| (Dahlhaus et al., 2016) | Cancer signaling in neuroblastoma | *Homo sapiens* |
| (Davila-Velderrain et al., 2015) | Early flower development | *Arabidopsis thaliana* |
| (Enciso et al., 2016) | Lineage fate decision of hematopoietic cells | *Homo sapiens* |
| (Fauré et al., 2006) | Mammalian cell cycle | *Homo sapiens* |
| (García-Gómez et al., 2017) | Root apical meristem | *Arabidopsis thaliana* |
| (Giacomantonio and Goodhill, 2010) | Cortical area development | *Homo sapiens* |
| (Gupta et al., 2007) | Neurotransmitter signaling | *Homo sapiens* |
| (Herrmann et al., 2012) | Cardiac development | *Homo sapiens* |
| (Irons, 2009) | Cell cycle | *Saccharomyces cerevisiae* |
| (Klamt et al., 2006) | T-cell receptor signaling | *Homo sapiens* |
| (Krumsiek et al., 2011) | Hematopoiesis | *Homo sapiens* |
| (MacLean and Studholme, 2010) | Type III secretion system | *Pseudomonas syringae* |
| (Mai and Liu, 2009) | Apoptosis | *Homo sapiens* |
| (Marques-Pita and Rocha, 2013) | Body segmentation | *Drosophila. Melanogaster* |
| (Martinez-Sanchez et al., 2015) | CD4+ T-cell fate | *Homo sapiens* |
| (Méndez and Mendoza, 2016) | B-cell differentiation | *Homo sapiens* |
| (Méndez-López et al., 2017) | Immortalization of epithelial cells | *Homo sapiens* |
| (Mendoza and Xenarios, 2006) | T-cell signaling | *Homo sapiens* |
| (Meyer et al., 2017) | Senescence-associated secretory phenotype | *Homo sapiens* |
| (Orlando et al., 2008) | Cell cycle | *Saccharomyces cerevisiae* |
| (Ortiz-Gutiérrez et al., n.d.) | Cell cycle | *Arabidopsis thaliana* |
| (Ríos et al., 2015) | Gonadal sex determination | *Homo sapiens* |
| (Saadatpour et al., 2011) | T-cell large granular lymphocyte survival | *Homo sapiens* |
| (Sahin et al., 2009) | Cell cycle | *Homo sapiens* |
| (Sankar et al., 2011) | Hormone crosstalk | *Arabidopsis thaliana* |
| (Siegle et al., 2018) | Aging of satellite cells | *Homo sapiens* |
| (Sridharan et al., 2012) | Oxidative stress | *Homo sapiens* |
| (Sun et al., 2014) | Endomesoderm tissue specification | *Sea urchin* |
| (Thakar et al., 2012) | Immune response | *Homo sapiens* |
| (Todd and Helikar, 2012) | Cell cycle | *Saccharomyces cerevisiae* |
| (Yousefi and Dougherty, 2013) | Metastatic melanoma | *Homo sapiens* |

### Test for scale-freeness

If a Boolean network has a scale-free network architecture, it can be described by the power law distribution *P*(*k*) ∝ *k*^−*α*^ where *α* is the power law scaling parameter. To identify scale freeness, we tested if the degree distribution of the network can plausibly be described by the power law distribution by using the R-package poweRlaw v.70.2 (Gillespie, 2015).

### Reducing network size

The size and the complexity of Boolean networks increases rapidly with each additional node and exhaustive search methods cannot be applied to larger networks. Therefore, we reduced the search space of large Boolean networks to accelerate the analysis. This was achieved by removing nodes which do not regulate other nodes. The procedure was repeated until all superfluous nodes were removed (Appendix 1 —Algorithm 3).

### Identify minimal set of nodes determining the dynamics

Two strategies were applied to determine a minimal set of compounds determining the dynamics of the complete network. The heuristic is defined in terms of significance. Here, the significance of a node is maximal if its transition function depends on its own value. Otherwise the significance of node *g* is equal to the number of nodes whose transition function depends on *g*. Therefore, in each iteration the heuristic selects a compound *g* with the highest significance until a set of implicant nodes is found (Appendix 1 —Algorithm 1). For reference we use an exhaustive search algorithm to find minimal implicant sets *G* of size *k* for increasing values of *k*. The algorithm terminates for the smallest value of *k* such that some subset *G*:*G* = *g*_*i*1_, …, *g_ik_* is an implicant set, i.e. for every assignment a network state is observable (Appendix 1— Algorithm 2).

### Partial assignments

A partial assignment defines the value of some nodes of the Boolean network. The value of the other nodes is undefined. The transition function 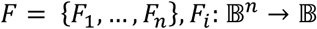 can be applied to a partial assignment as follows: If the value of the transition function *F_i_* of a node *i* is uniquely determined from nodes which are defined under the partial assignment then the node is assigned this value. The value of all other nodes remains unchanged. Repeated application of the transition function to a partial assignment will define the value of not necessarily all nodes. If the assignment can be extended to all nodes and hence defines a state of the network, we say that the network state is observable from the partial assignment. A set of nodes is called implicant if for every assignment to the nodes a network state is observable. An implicant is minimal if the cardinality of the set is minimal.

### Connectivity

The static connectivity of nodes within a Boolean network is defined by their incoming and outgoing edges. To identify static hub nodes, we calculated the z-score: 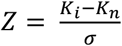 whereby *K_i_* is the number of interactions of component *i* to other components in the network, *K_n_* is the average of interactions of all components within the network and *σ* is the standard deviation of *K_n_*. Thereby, components with a z-score >2.5 are considered as hub nodes (Guimerà and Nunes Amaral, 2005). In addition to the static interactions deriving from the connections of nodes within the network, we also considered the biological interactions of each protein included in the Boolean network. Therefore, protein interaction tables of each organism considered in one of our analyzed Boolean network models were obtained from BioGRID (Stark et al., 2006) (Version 3.5.168) and the directed interactions of each protein were counted with R (R Core Team, n.d.). In the case of Boolean network nodes consisting of several proteins or nodes whose proteins can be several isoforms, the average of all these possible proteins was taken. Afterwards, the z-score was used again to define biological hubs.

### Statistical analyses

Analyses and visualization were done with R (R Core Team, n.d.) (https://www.r-project.org). All statistical tests are two-sided.

Figure 2B/C: Statistics were performed with Cochran’s Q test and post-hoc pairwise sign test with Bonferroni correction (R package RVAideMemoire (Hervé, n.d.)).

Figure 2D/E: Effects of interventions were analyzed by Wilcoxon test (implicants vs hubs/other nodes).

Figure 3B/C/D: Experimental data were analyzed by Wilcoxon test (each treatment vs untreated).

### Model establishment of MAPK/Wnt crosstalk in colorectal cancer

The model was constructed based on literature search and interactions taken from the curated databases BioGRID (Stark et al., 2006) and Metacore™ (Thomson Reuters Inc., Carlsbad, CA). For the model setup, key components of MAPK-signaling and canonical Wnt-signaling cascade and their crosstalk components were considered. A short model description with all Boolean functions and their references can be found in Appendix 2—Table 1.

### Cell line

SW480 cells were obtained from the American Type Culture Collection (ATCC). Cells were cultured at 37°C and 5% (v/v) CO_2_ in DMEM high glucose (Sigma-Aldrich) medium containing 10% fetal bovine serum (Life technologies) and 1% Penicillin-Streptomycin (Life technologies). Cells were routinely tested for absence of mycoplasma (GATC).

### Drug treatment

Drug treatments were performed in 6-well plates 24 hours after seeding, cells were treated with 1 mL of medium containing 2 μM BVD-523 (HY-15816, MedChemExpress) or 12 μM TD-52 (SML2145, Sigma-Aldrich) or a combination of BVD-523 and TD-52. Drugs were dissolved in DMSO (Sigma-Aldrich).

### Cell count/ Proliferation assay/ Apoptosis measurement

10 μL of cells were mixed with Trypan blue (Life technologies), loaded on a cell counting slide (Countess™, Invitrogen) and counted with the Counter II (Invitrogen). Here, we also detected the amount of living viable and dead cells.

### Wound healing assay

Cells were seeded on fibronectin coated cover slips. After cells were confluent, they were treated as described before. The wound was introduced with a 200 μL pipette tip. Series of pictures of each wound were taken (BIOREVO BZ-9000, KEYENCE, magnification 4x) and merged (program BZ analyser II, KEYENCE) to a complete picture of the wound that was analyzed further. Wound closing was measured by calculating whole wound border areas at different time points for each repetition with the “MRI wound healing tool” implemented in Image-J.

### Immunofluorescence staining

SW480 cells were grown on glass coverslips. Cells were fixed with 4% PFA for 15 min and were permeabilized with 0.1% Triton X-100 for 10 min. Cells were blocked in 0.5% BSA for 45 min, incubated with primary (E-cadherin #610181, BD Bioscience, 1:200) and secondary antibodies (anti-mouse Alexa 488, Dianova, 1:1000) for 2 h and 1h at RT each, and were mounted in DAPI mounting medium. Images were taken with a Leica TCS SP5 II confocal microscope in a single plane (63-objective) using LAS AF software. Same exposure settings were used in controls and drug treated samples. Images of the same experiment were equally processed using Adobe Photoshop CS6 software.

### Checking for essentiality

All genes of the human networks were screened for essentiality using the HEGIAP database (Chen et al., 2019). Here, only essential genes for homo sapiens were considered.

### Code availability

The pseudocode for our analyses is included in the Appendix 1. The complete code for our simulations is available upon request.

## Acknowledgments

This work was supported by SFB 1074 (DFG) German Science Foundation (DFG, grant number 217328187), German Research Foundation GRK 2254 HEIST, Federal Ministry of Education and Research (BMBF, TRANSCAN VI – PMTR-pNET, ID 01KT1901B).

